# MetaFlow: Metagenomic profiling based on whole-genome coverage analysis with min-cost flows

**DOI:** 10.1101/038208

**Authors:** Ahmed Sobih, Alexandru I. Tomescu, Veli Mäkinen

## Abstract

High-throughput sequencing (HTS) of metagenomes is proving essential in understanding the environment and diseases. State-of-the-art methods for discovering the species and their abundances in an HTS metagenomic sample are based on genome-specific markers, which can lead to skewed results, especially at species level. We present MetaFlow, the first method based on coverage analysis across entire genomes that also scales to HTS samples. We formulated this problem as an NP-hard matching problem in a bipartite graph, which we solved in practice by min-cost flows. On synthetic data sets of varying complexity and similarity, MetaFlow is more precise and sensitive than popular tools such as MetaPhlAn, mOTU, GSMer and BLAST, and its abundance estimations at species level are two to four times better in terms of *ℓ*_1_-norm. On a real human stool data set, MetaFlow identifies *B.uniformis* as most predominant, in line with previous human gut studies, whereas marker-based methods report it as rare. MetaFlow is freely available at http://cs.helsinki.fi/gsa/metaflow

## 1 Introduction

Microbes—microscopic organisms that cannot be seen by the eye, which include bacteria, archaea, some fungi, protists and viruses—are found almost everywhere, in the human body, air, water and soil, and play a vital role in maintaining the balance of ecosystems. For example, diazotrophs are solely responsible for the nitrogen fixation process on earth [17], and half of the oxygen on earth is produced by marine microbes [19].

Metagenomic sequencing allows studying microbes sampled directly from the environment without prior culture. A fundamental analysis of a metagenomic sample is finding what species it contains (the sample richness) and what are their relative abundances. This is a challenging task due to the similarity between the species' genomes, sequencing errors, and the incompleteness of reference microbial databases.

One of the first methods, applicable only at high taxonomic levels [15], focused on 16S ribosomal RNA marker genes. Its limitations have been mitigated by high-throughput sequencing of the entire genomes in a metagenomic sample, and a number of methods dealing with this data have been proposed. The simplest of them is to align the reads to a reference database, using e.g. BLAST [1], and to choose the best alignment for every read. This approach cannot break the tie between multiple equally-good alignments of a read, and it cannot detect false positive alignments. One way of avoiding alignment ties, but not false positive alignments, is to estimate the sample structure only at a high taxonomic level. For example, MEGAN [7] assigns each read with ties to the lowest common ancestor of its alignments in the reference taxonomic tree. Another method for breaking ties was proposed in PhymmBL [2], based on Interpolated Markov Models (IMMs). For each read, it combines the BLAST alignment score to a particular genome, with another score based on the probability of the read being generated by an IMM of that genome. This results in a single maximum-scoring alignment for every read, which improves over basic BLAST alignments. This method still cannot eliminate false alignments, and does not scale to high-throughput sequencing samples. For example, it takes more than one hour to classify 5,730 reads of length 100bp [2].

The current state-of-the-art approach is to construct a small curated reference database of genomic markers. These markers can be clade-specific marker genes (MetaPhlAn [20]), universal marker genes (mOTU [22]), or genome-specific markers not restricted to coding regions (GSMer [23]). The reads are aligned only to these marker regions, which makes it a fast process, and the estimations are more accurate at the species level because of the marker curation process. However, due to the short length of the marker regions, the abundance estimations can be extremely skewed in some cases. In addition, the markers are uniquely identifying the microbial genomes only among the currently known genomes, meaning that the entire database of markers must be re-computed with the addition of each new reference microbial genome.

In this paper, we propose a new method which addresses the problems of equally-good alignments and of false alignments, with accurate predictions at the species level. Our method takes into account the entire read alignment landscape inside all genomes in the reference database. The main idea is to exploit the assumption that if enough reads are sampled, then these reads will cover most of the genomic regions. In addition, not relying on specific genomic markers, there is no need to curate a reference database. Our problem formulation is based on a matching problem applied to a bipartite graph constructed from read alignments. This problem is NP-hard, but we give a practical strategy for solving it with min-cost network flows. See Fig. 1 for an overview of all methods.

**Fig. 1.**
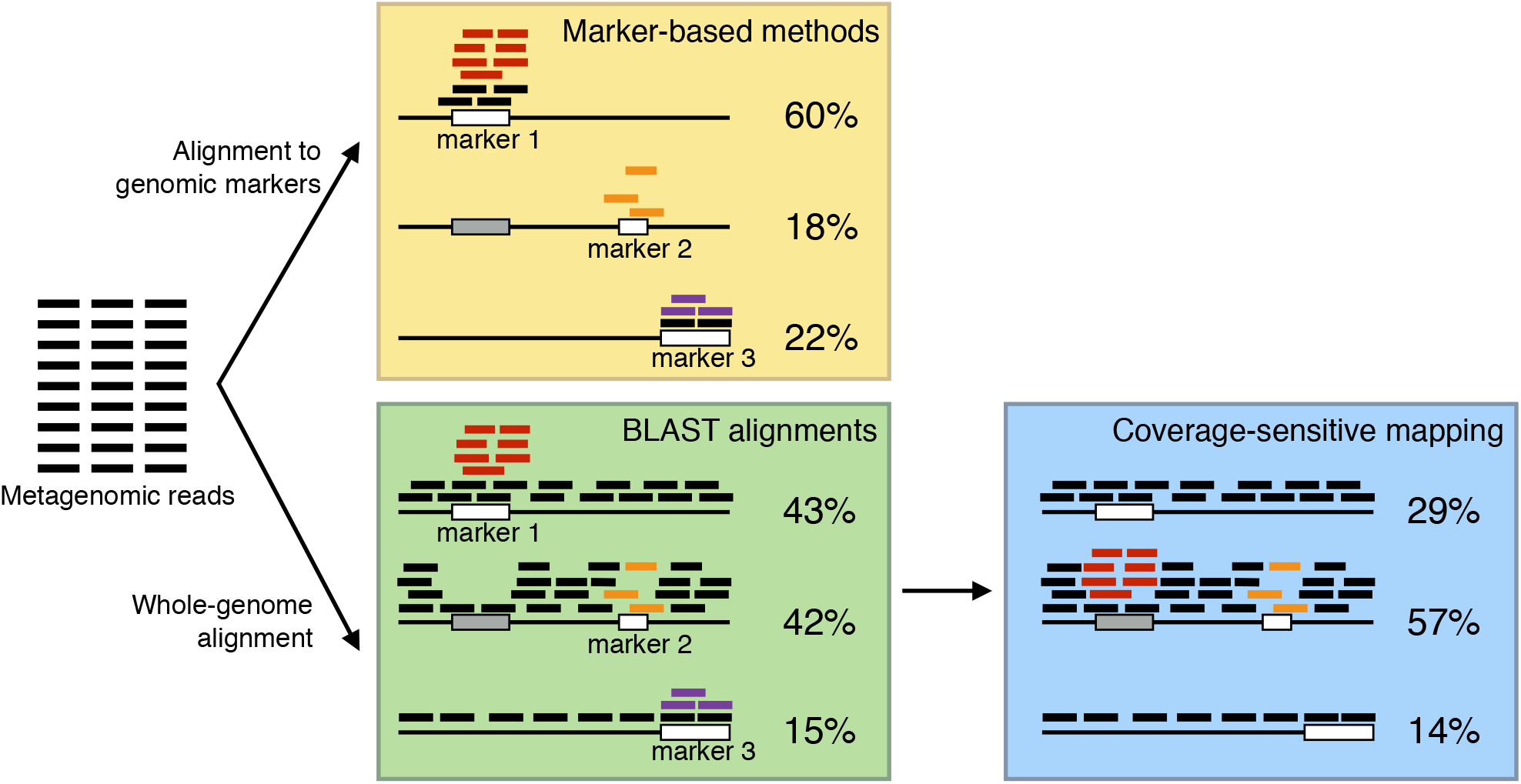
Overview of the methods compared in this paper. In the yellow box, reads are aligned only to genomic markers, and the relative abundances are highly skewed: marker 1 receives more false mappings (in red) because it is similar with a subsequence of the second genome (gray); marker 2 has a drop in coverage due to a sequencing bias, and it is covered only by three reads (orange); marker 3 is covered also by some reads sequenced from a species not in the reference database (violet). In the green box, reads have whole-genome BLAST alignments, but the relative abundances are still skewed: the tie in the alignment of the red reads is not resolved, and the violet reads from an unknown species aligning to the third genome are not removed. In the blue box, the coverage-sensitive mapping of the reads: the red reads are correctly aligned to the second genome (in the gray sequence) and the violet reads from an unknown species are discarded from the third genome. The abundances are relative to the known genomes.

We performed experiments on synthetic and real data sets, and compared MetaFlow with popular tools MetaPhlAn [20], mOTU [22], GSMer [23], and standard BLAST alignments. MetaFlow outperforms the other methods in its ability to correctly identify the species and their abundance levels. On synthetic data, MetaFlow’s predictions are more precise and sensitive, and its abundance estimations are up to two to four times better in terms of *ℓ*_1_-norm. On a real fecal metagenomic sample from the Human Microbiome Project, MetaFlow reports *B.uniformis* as most predominant species, in line with previous human gut studies [16]. However, marker-based methods MetaPhlAn and mOTU assign a very low abundance to it.

## 2 Methods

We assume in input a set of BLAST hits of the metagenomic reads inside a collection of reference genomes, which we call *known genomes*. The output is the richness of the sample and the relative abundance of each known species. This is obtained by selecting, for every read, exactly one hit in a reference genome, or classifying the read as originating from a species not in the reference database (we call such species *unknown*).

The optimal selection is the one simultaneously achieving the following three objectives: (1) few coverage gaps, (2) uniform coverage, and (3) agreement between BLAST scores and final mappings. Objective (1) allows the detection of outlier genomes that have only few regions covered. Objective (2) breaks ties between read alignments, and is also based on detecting abnormal read coverage patterns.

We model the above-mentioned input for this problem as a bipartite graph, such that the reads form one part of the bipartition, and the reference genomes form the other part of the bipartition. Objectives (1)-(3) are not modeled independently, but combined in a single objective function, as discussed in the next section. Our modeling is inspired by the interesting Coverage-sensitive many-to-one min-cost bipartite matching problem, introduced in [11] for mapping reads to complex regions of a reference genome. We extended this model to our metagenomic context, since reads can have mappings to more than one reference genome, or can originate from unknown species.

### 2.1 Problem formulation and computational complexity

Assume that the reads have BLAST hits in the collection of reference genomes 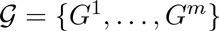. We partition every genome *G^i^* into substrings of a fixed length *L*, which we call *chunks*. Denote by *s_i_* the number of chunks that each genome *G^i^* is partitioned into. We construct a bipartite graph *G* = (*A* ∪ *B, E*), such that the vertices of *A* correspond to reads, and the vertices of *B* correspond to the chunks of all genomes *G*^1^,…,*G^m^*. More specifically, for every chunk *j* of genome *G^i^*, we introduce a vertex 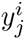, and we add an edge between a read *x* ∈ *A* and chunk 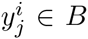 if there is a BLAST mapping of read *x* starting inside chunk *j* of genome *G^i^*. This edge is assigned the cost of the mapping (BLAST scores can be trivially transformed to costs), which we denote here by 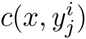. In order to model the fact that reads can originate from unknown species (whose genome is not present in the collection 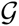), we introduce an ‘unknown’ vertex *z* in *B*, with edges from every read *x* ∈ *A* to *z*, and with a fixed cost *c*(*x, z*) = *γ*, where *γ* is appropriately initialized.

In the *coverage-sensitive metagenomic mapping* problem stated below, the tasks are: first, for each *G^i^*, to find the number of reads sequenced (i.e., originating) from it, which we denote by *r_i_* (*r_i_* is 0 if *G^i^* is an outlier); second, to select an optimal subset *M* ⊆ *E* such that for every read *x* ∈ *A* there is exactly one edge in *M* covering it (i.e., a mapping of *x*). These must minimize the sum of the following two costs:

A. the sum, over all chunks of every genome *G^i^*, of the absolute difference between its due read coverage, namely *r_i_/s_i_*, and the number of read mappings it receives from *M* (this corresponds to Objective (2));
B. the sum of all edge costs in *M* (this corresponds to Objective (3)).

Our formal problem definition is given below. We use the following notation: *n* is the number of reads, *m* is the number of different genomes where the reads have BLAST hits, *s_i_* is the number of chunks of each genome *G^i^, i* ∈ {1,…, *m*}. In a graph *G* = (*V, E*), *d_M_*(*v*) denotes the number of edges of a set *M ⊆ E* incident to a vertex *v*.

##### Coverage sensitive metagenomic mapping

**Input:**

– a bipartite graph *G* = (*A* ∪ *B, E*), where *A* is the set of *n* reads, *B* = 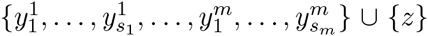 is the set of all genome chunks plus *z*, the ‘dummy’ node,
– a cost function *c: E* → ℚ,
– constants *α* ∈ (0,1), *β*, *γ* ∈ ℚ_+_.

**Tasks:**

– find a vector *R* = (*r*_1;_…, *r_m_*) containing the number of reads sequenced (i.e., originating) from each genome *G^i^, i* ∈ {1,…, *m*}, and
– find a subset *M* ⊆ *E* such that *d_M_*(*x*) = 1 holds for every *x* ∈ *A* (i.e., each read is covered by exactly one edge of *M*),

which together minimize:

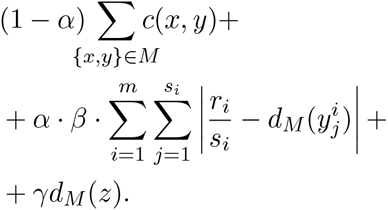

We first show that this problem is computationally intractable.

##### Theorem 1. The coverage-sensitive metagenomic mapping problem is NP-hard.

*Proof*. We reduce from the Exact cover with 3-sets (X3C) problem (see e.g., [5]). In this problem, we are given a collection S of 3-element subsets *S*_1_,…,*S_m_* of a set *U* = {1,…,*n*}, and we are required to decide if there is a subset 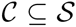, such that every element of *U* belongs to exactly one 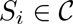.

Given an instance of Problem X3C, we construct the bipartite graph *G* = (*A* ∪ *B, E*), where *A* = *U* = {1,…, *n*}, and *B* corresponds to *S* in the following sense:

– *s_i_* = 3, for every *i* ∈ {1,…, *m*}
– for every *S_j_* = {*i*_1_ < *i*_2_ < *i*_3_}, we add to *B* the three vertices *y*_*j*,1_, *y*_*j*,2_, *y*_*j*,3_ and the edges {*i*_1_,*y*_*j*,1_}, {*i*_2_,*y*_*j*,2_}, {*i*_3_,*y*_*j*,3_}, each with cost 0.

For completeness, we also add vertex *z* to *B*, and edges of some positive cost between it and every vertex of *A*.

We now show that an instance for Problem X3C is a ‘yes’ instance if and only if the coverage-sensitive metagenomic mapping problem admits on this input a solution set *M* ⊆ *E* of cost 0. Observe that, in any solution *M* of cost 0, the numbers *r_i_* of reads sequenced from each genome are either 0 or 3, and vertex *z* has no incident edges in *M*.

For the forward implication, let 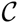 be an exact cover with 3-sets. We choose the vector *R* = (*r*_1_,…, *r_m_*) as follows:

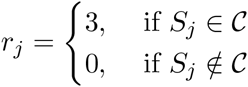

and we construct *M* as containing, for every *S_j_* = {*i*_1_ < *i*_2_ < *i*_3_} ∈ 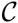, the three edges {*i*_1_, *y*_*j*,1_}, {*i*_2_, *y*_*j*,2_}, {*i*_3_, *y*_*j*,3_}. Clearly, these *R* and *M* give a solution of cost 0 to the coverage-sensitive metagenomic mapping problem.

Vice versa, if the coverage-sensitive metagenomic mapping problem admits a solution (*R, M*) of cost 0, thus, one in which the numbers *r_i_* of reads sequenced from each genome are either 0 or 3, and *z* has no incident edges, then we can construct an exact cover with 3-sets 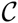 by taking:

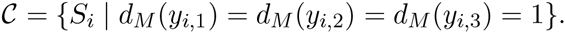

Due to the hardness result from Thm. 1, we opt for the common *iterative refinement strategy*, akin to the strategy behind *k*-means clustering [21], or Viterbi training strategies with Hidden Markov Models [4]. In Sec. 2.3 we present the details of this approach, but the main ideas are:

1. If the unknown vector *R* = (*r*_1_,…, *r*_m_) is fixed to some value *R* = (*a*_1_,…, *a_m_*), then finding only the optimal mapping *M* can be solved in polynomial-time with min-cost flows. We show this in Sec. 2.2.
2. For finding the optimal *R* (and *M*), we start with a vector *R*^0^ = (*a*_1_,…, *a_m_*), where *a_i_* equals the number of reads with BLAST hits to *G^i^*. We repeat the following process, until the vector *R* converges to a stable value. For each iteration *j*:

a. compute the optimal read mapping *M^j^* by min-cost flows (as we will show in Sec. 2.2), using *R^j^* as input;
b. update *R^j^* to *R*^*j*+1^, a vector whose *i*-th component equals *mean_i_ · s_i_*; here *mean_i_* is the 20%-trimmed mean read coverage of the chunks of genome *G^i^*, obtained from *M^j^*.

### 2.2 The reduction to a min-cost flow problem

We refer the reader to Sec. A.1 of the Supplementary Material for some standard notions and definitions of flow networks and min-cost flows. Given an input bipartite graph *G* = (*A* ∪ *B, E*) for the coverage-sensitive metagenomic mapping problem, with a fixed vector *R^j^* = (*a*_1_,…, *a_m_*) containing the number of read sequenced from the *m* genomes (at an iteration *j*), we construct a flow network *N* = (*G*, ℓ, u, c, q*) (see Fig. 2(b)). Here, *ℓ, u*: *E*(*G*\*) → ℕ are the *demand* and *capacity* of every arc, respectively; *c*: *E(G*)* → ℚ is the function giving the cost per unit of flow for every arc of *G*\*; *q* is the required value of the flow, that is, the value of the flow that must exit the unique source of *G*^*^, which equals the value of the flow that must enter the unique sink of *G*^*^.

**Fig. 2.**
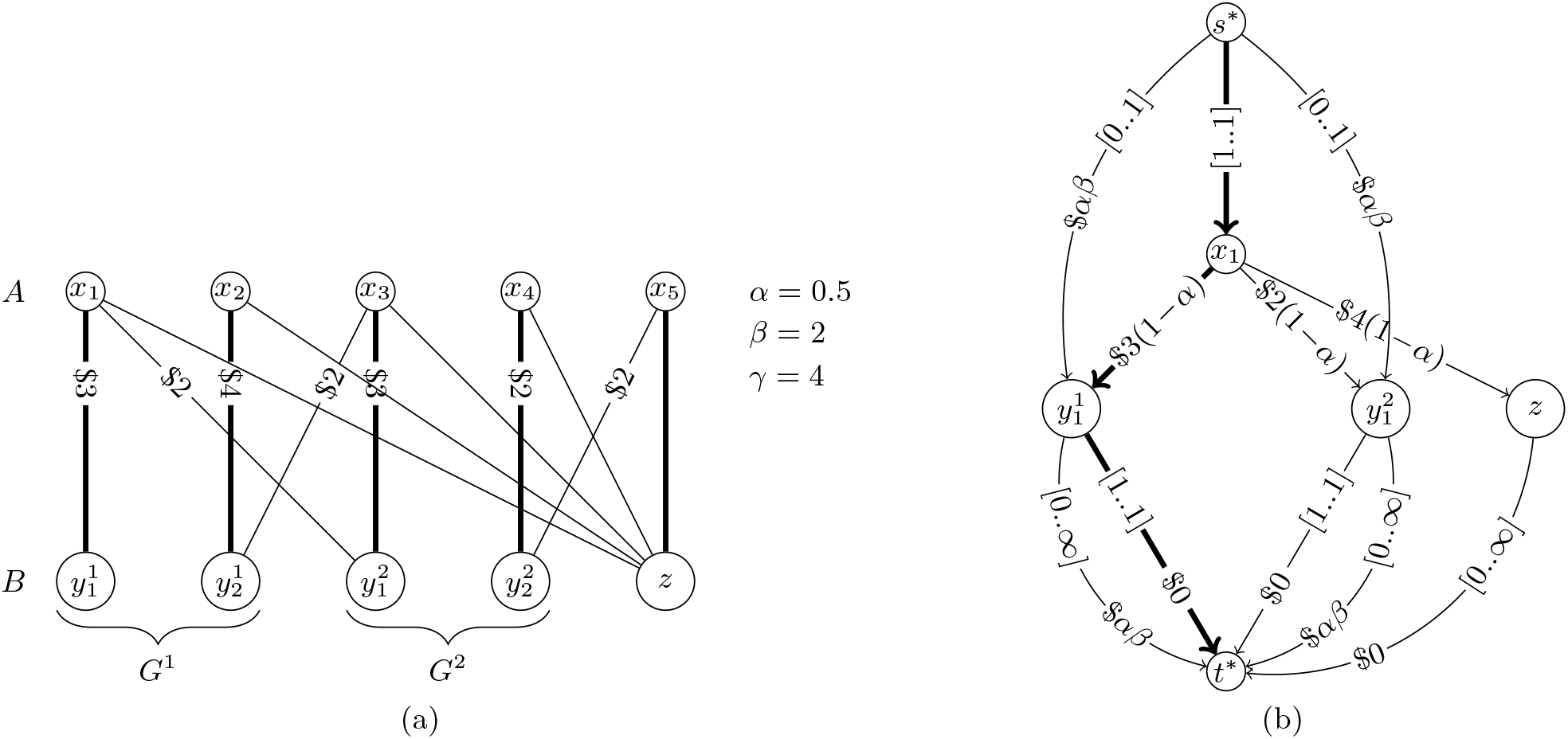
Fig. 2(a) A bipartite graph *G* = (*A ∪ B, E*), whose edges are labeled with costs. The costs of the edges incident to *z* are not drawn, and they all equal *γ*. An optimal solution for the coverage-sensitive metagenomic mapping is shown (highlighted edges). The optimal read mappings to *G*^1^ and *G*^2^ are *r*_1_ = *r*_2_ = 2, and the optimal value of the objective function is (1 - *α*)12 + *αβ*((1 - 1 + 1 - 1) + (1 - 1 + 1 - 1)) + *γ* = 10. Fig. 2(b): A subpart of the flow network *N* constructed from *G*, corresponding to the vertices 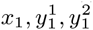. The arcs with flow value 1 arc highlighted. The demands and capacities of an arc (*u, v*) are indicated as [*ℓ(u, v)..u(u, v)*] and its cost as $*c*(*u, v*).

The construction of *G*\* starts with *G*, and orients all its edges from *A* towards *B*, and sets their demand to 0, their capacity to 1, and their cost as in the input metagenomic mapping instance, multiplied by (1 – *α*). We add a global source *s*\* with arcs toward all vertices in *A*, with demand and capacity 1, and 0 cost. Since we pay the fixed penalty *α · β* for every read below or above the given coverage of each genome chunk 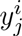, we add the following arcs:

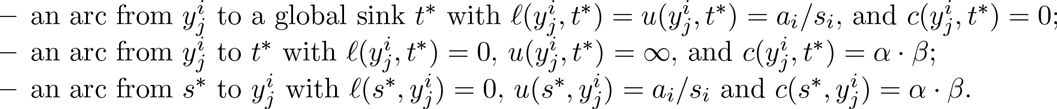

We also add arcs from every vertex *x ∈ A* to the unknown vertex *z ∈ B*, with demand *ℓ(x, z)* = 0, *u(x,z)* = 1, and cost *c(x,z)* = *γ*. From *z* we add an arc to *t*\* with *ℓ*(*z, t*\*) = 0, *u*(*z, t*\*) = ∞, *c*(*z, t*^*^) = 0.

Finally, we set 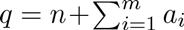, and also add the arc (*s*\*, *t*\*) with 0 demand and cost, and infinite capacity. The optimal matching for the coverage-sensitive metagenomic mapping problem consists of the edges of *G* whose corresponding arcs in *G*^*^ have non-zero flow value in the integer-valued min-cost flow over *N*. See Fig. 2 for an example.

The correctness of this reduction can be seen as follows. As long as a genome chunk 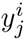 has exactly *a_i_*/*s_i_* reads mapped to it, then no cost is incurred; this is represented by the arc 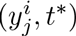 with demand and capacity *a_i_*/*s_i_* and 0 cost. If 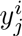 receives more reads than *a_i_/s_i_* then these additional reads will flow along the parallel arc 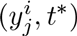 of cost *α · β*. If 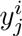 receives less reads than *a_i_/s_i_* then some compensatory flow comes from *s*\* through the arc 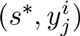, where these fictitious reads again incur cost *α · β*. We set the capacity of 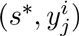 to *a_i_/s_i_* since at most *a_i_/s_i_* fictitious reads are required to account for the lack of reads for genome chunk 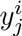.

Requiring flow value 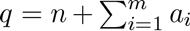 ensures that there is enough compensatory flow to satisfy the demands of the arcs of type 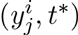. Having added the arc (*s*\*, *t*\*) with 0 cost ensures that there is a way for the exogenous flow which is not needed to satisfy any demand constraint to go to *t*\*, without incurring an additional cost.

### 2.3 Overview of the implementation

Our practical implementation is divided into five stages. These depend on some parameters, whose complete list we give in Sec. B.3 of the Supplementary Material.

**Stage 1: Removing outliers species** A genome *G^i^* ∈ 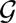 is considered an outlier if at least one of the following conditions holds.

– The average read coverage of *G^i^* (i.e., the number of reads with BLAST hits to *G^i^* multiplied by the average read length and divided by the length of *G^i^*) is lower than a given parameter.
– The average read mapping per chunk (i.e., the number of reads with BLAST hits to *G^i^* divided by *s_i_*) is lower than a given parameter.
– The percentage of chunks without any BLAST hit is more than a given parameter.

In this stage, we remove each outlier genome *G^i^* and the reads that have BLAST hits only to *G^i^*.

**Stage 2: Breaking ties inside each genome** A read can have BLAST hits to different chunks of the same genome. In this stage, for each read remaining after Stage 1, we select only one BLAST hit in each genome, as follows. For each remaining genome 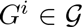, we create a sub-problem instance *G* = (*A* ∪ *B, E*) where *A* consists only of the reads that have BLAST hits to *G^i^*, and *B* consists only of the chunks of *G^i^* (excluding the unknown vertex *z*, which will be dealt with in Stage 4). We fix the one-element vector *R* as *R* = (*r*_1_) = (|*A*|), and solve this sub-problem using the min-cost flow reduction described in Sec. 2.2. After this stage, every read has at most one hit to each genome, but it can still have hits to multiple genomes.

**Stage 3: Breaking ties across all genomes** A read can be mapped to different species, due to the similarity between their genomes. In order to select only one read mapping across all genomes, we solve the coverage-sensitive metagenomic mapping problem on a graph *G* = (*A* ∪ *B, E*), as follows. The set *A* consists of all remaining reads, and the set *B* of all chunks of the remaining genomes. The set of edges *E* is the one obtained after the filtration done in Stage 2. Since this problem is NP-hard, we employ the iterative refinement strategy, coupled with min-cost flows, mentioned at the end of Sec. 2.1. After each iteration *j*, we use the resulting mapping *M^j^* to remove outlier genomes, as in Stage 1. After this stage, each read is mapped to exactly one genome, and to only one of its chunks.

**Stage 4: Identifying reads from unknown genomes** In this stage we identify reads originating from species whose reference genomes are not present in the reference database. We run the same min-cost flow reduction as in Stage 2, to which we add the unknown vertex *z*. If a read is mapped to *z*, then it will be marked as coming from an unknown genome and removed from the graph. Finally, we again remove outlier genomes, as in Stage 1.

**Stage 5: Estimating richness and relative abundances** For every genome *G^i^*, we compute its average read coverage *read_coverage*(*i*), and its relative abundance *rel_abundance*(*i*) as: *read_coverage*(*i*) = 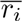 · *R/length(i), rel_abundance(i)* = *read_coverage*(*i*)/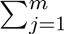*read_coverage*(*j*), where 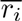 is the number of reads mapping to *G^i^* after Stages 1-4, *R* is the average read length, and *length*(*i*) is the length of *G^i^*.

## 3 Experimental setup

Our experimental setup measures the effect of the following three factors: (1) genotypic homogeneity between the species, (2) complexity of the sample (i.e. the number of species), and (3) the presence of species unknown to the methods. We simulated two types of data sets: a *low complexity* type (LC), consisting of 4M reads sampled from 15 different species, and a *high-complexity* type (HC), consisting of 40M reads sampled from 100 different species. The goal of the simulated LC data sets is to evaluate the methods under different levels of genotypic homogeneity between the species. For example, species with high genotypic homogeneity are difficult to precisely identify and their presence usually leads to incorrect predictions. We explain these levels of similarity in Table 4 of the Supplementary Material. The simulated HC data sets test how a large number of randomly chosen species influences the accuracy of the methods. As opposed to the LC data sets, they do not test the ability of the tools to deal with similar species. This experimental setup is in line with previous studies [20,13].

In both the LC and HC data sets, we had two experimental scenarios: one in which all species in the sample are known to the methods (LC-Known, HC-Known), and one in which a fraction of them are unknown (LC-Unknown, HC-Unknown). LC-Known is the “perfect-information” scenario, which though not realistic, shows the performance of the tools in the best possible conditions. HC-Unknown is the most realistic scenario. In total, these simulated experiments contain 48 data sets.

The abundances of the species in each data set were chosen log-normal distributed (with mean=1, standard deviation=1), also in line with previous experiments [20]. This presents a challenge in finding less abundant species, since the ratio between the most abundant species and the least abundant is 100 in most cases, and the top 10% most abundant species represents about 35% of the sample. We selected in total 817 bacterial species from the NCBI microbial genome database, and used Metasim [18] to create the data sets using the default 454-pyrosequencing error model (with mean read length=250 and standard deviation=1).

In order to evaluate the accuracy of the richness estimations, we evaluated the *sensitivity* (also called *recall*) and the *precision*. Sensitivity is defined as *TP/NS*, and precision is defined as *TP/(TP + FP)*, where *TP* is the number of species correctly identified by the tool, *FP* is the number of species not present in the sample but reported by the tool, and *NS* is the true number of species in the sample. In order to obtain a single measure of accuracy, we also computed the harmonic mean of precision and specificity, known as *F-measure*, and defined as 2 · *precision · sensitivity/(precision + sensitivity)*. To evaluate the accuracy of the relative abundance predictions, we measured the *ℓ*_1_-norm of these abundances for each data set, expressed in percentage points. For the data sets with unknown genomes, we excluded the unknown genomes from the abundance vectors and we re-normalized the resulting relative abundances to the known genomes. Note that some methods, such as MetaPhlAn [20] and mOTU [22], give some abundance estimations for the unknown genomes at a higher taxonomic level (e.g. genus). Our method is focused instead on the core problem of analyzing the known species, without attempting to estimate the relative abundances of the unknown genomes.

**Table 1.** Average over the F-measure and *ℓ*_1_-norm in each experimental scenario. The 15 LC data sets contain 4M reads from 15 species, and the 9 HC data sets contain 40M reads from 100 species.

|  | LC-Known |  | LC-Unknown |  | HC-Known |  | HC-Unknown |  |
| --- | --- | --- | --- | --- | --- | --- | --- | --- |
| | F-measure | $\ell_1$-norm | F-measure | $\ell_1$-norm | F-measure | $\ell_1$-norm | F-measure | $\ell_1$-norm |
| MetaFlow | **0.971** | **10.41** | **0.825** | **17.87** | **0.976** | **4.86** | **0.883** | **8.01** |
| MetaPhlAn | 0.946 | 26.42 | 0.770 | 31.48 | 0.958 | 18.61 | 0.844 | 19.13 |
| BLAST | 0.909 | 12.46 | 0.745 | 21.47 | 0.920 | 5.94 | 0.809 | 11.25 |
| GSMer | 0.218 | N/A | 0.163 | N/A | 0.327 | N/A | 0.259 | N/A |
| mOTU | 0.924 | 36.31 | 0.780 | 43.74 | 0.949 | 10.55 | 0.847 | 18.73 |

We compared the performance of MetaFlow against BLAST [1], MetaPhlAn [20], mOTU [22] and GSMer [23]. In the BLAST analysis, we always selected the best alignment; in case of multiple equally-good alignments, we randomly selected one of them; if the read coverage of a species is below 0.3 X, then we considered it an outlier. GSMer does not provide relative abundances, so we compared only the accuracy of the richness estimations. For mOTU, some species among our known genomes were not covered in its database, so in our evaluation it received full marks on these species. On the other hand, some of the species chosen as unkown in our experiments already existed in mOTU’s database. For these species, we removed mOTU’s correct prediction.

We also ran our tool using a real data set. We merged 6 G_DNA_Stool samples of a female from Human Microbiome Project (5 samples were generated using Illumina, and one sample using LS454). Their accession numbers are in the Supplementary Material. The read length of all reads was normalized to 100bp. The total number of reads from all samples was 287,565,377, out of which 98,223,162 BLAST mapped to one or more species. Only alignments with identity ≥ 97% were selected as an input for MetaFlow. In addition to the full reference genomes in NCBI’s microbial database, we also used two other references: a supercontig of *B.uniformis* (accession number NZ_JH724260.1), because *B.uniformis* was previously reported as most abundant in fecal samples [16]; and the longest scaffold of *B.plebeius* (accession number NZ_DS990130.1) because MetaPhlAn and mOTU report *B.plebeius* as most abundant in this sample.

See Table 2 for the running times of the methods tested in this paper.

**Table 2.** Running times of the methods as data set size increases. MetaFlow starts the analysis from the BLAST alignments (column “BLAST”). The data sets of size 4M and 40M are the synthetic ones; the reported running times are averages over all data sets of the same size. The data set of size 280M is the real one.

| Data set size | BLAST | MetaFlow | MetaPhlAn | mOTU | GSMer |
| --- | --- | --- | --- | --- | --- |
| 4M | 243 min | 28 min | 14 min | 9 min | 42 min |
| 40M | 1572 min | 459 min | 132 min | 84 min | 364 min |
| 280M | 3696 min | 2025 min | 387 min | 380 min | N/A |

## 4 Discussion

### 4.1 Synthetic data sets

We summarize the experimental results on simulated data in Table 1 and in Fig. 3. The complete results on each LC data set are in Tables 5 and 6 of the Supplementary Material. Since BLAST reports alignments in all the reference genomes, its sensitivity is always the maximum achievable. However, this comes at the cost of low precision, since there is no proper strategy for breaking ties among alignments, or for properly removing outlier genomes. MetaPhlAn and mOTU have better precision and F-measure than BLAST, confirming that marker genes are a good way of distinguishing between similar species. However, the sensitivity of the marker-based methods suggests that such an approach is not always accurate in identifying all species present in the sample. For example, MetaPhlAn’s sensitivity is not maximum in the LC12,LC13,LC14-Known data sets, even though these are perfect-information scenarios.

**Table 3.** Output files

| File | Description |
| --- | --- |
| abundance.csv | The main output file. It contains the final estimation of the species richness and abundance. |
| dist.csv | It contains the final distribution of the reads over all chunks in all genomes. |
| stage0.abundance.csv | Intermediary internal files that contain the estimation of the species richness and abundance before starting and after the stages 1,2, and 3, respectively. |
| stage1.abundance.csv |  |
| stage2.abundance.csv |  |
| stage3.abundance.csv |  |
| stage0.dist.csv | Intermediary internal files that contains the distribution of the reads over the chunks before starting and after the stages 1,2, and 3, respectively. |
| stage1.dist.csv |  |
| stage2.dist.csv |  |
| stage3.dist.csv |  |
| .log | The running log file. |

**Table 4.** Details of the LC-Known data sets. Each data set contains 4M reads from 15 species. The similarity level was determined by measuring the genomic-distance index with Mummer [9]

| data set name | Similarity level | Description |
| --- | --- | --- |
| LC1, LC2, LC3, LC4 | Very low-similarity | Species selected from 15 different genera, selected at random |
| LC5, LC6, LC7, LC8 | Very low-similarity | Species selected at random (there might be more from the same genus) |
| LC9 | Low-similarity | One species from the Lactobacillus genus, the rest randomly selected |
| LC10 | Low-similarity | Two species from Lactobacillus genus, the rest randomly selected |
| LC11 | Medium-similarity | One species from Shigella genus, the rest randomly selected |
| LC12 | Medium-similarity | Three species from Shigella genus, the rest randomly selected |
| LC13 | High-similarity | One species from Brucella genus, the rest randomly selected |
| LC14 | High-similarity | Three species from Brucella genus, the rest randomly selected |
| LC15 | High-similarity | Species randomly selected from only: Streptococcus, Brucella and Bacteroides |

**Table 5.** Detailed results on the LC-Known data sets. Sample descriptions are in Table 4. Best results are shown in bold. TP = true positives, FP = false positives, FN = false negatives.

| Sample ID | Tool | TP | FP | FN | Sensitivity | Precision | F-measure | $\hat{f}_1$-norm |
| --- | --- | --- | --- | --- | --- | --- | --- | --- |
| LC1 | MetaFlow | 15 | 0 | 0 | 1.000 | 1.000 | **1.000** | 0.72 |
|  | MetaPhlAn | 15 | 1 | 0 | 1.000 | 0.938 | 0.968 | 5.01 |
|  | BLAST | 15 | 0 | 0 | 1.000 | 1.000 | **1.000** | **0.44** |
|  | GSMer | 7 | 31 | 8 | 0.467 | 0.184 | 0.264 | N/A |
|  | mOTU | 15 | 1 | 0 | 1.000 | 0.938 | 0.968 | 7.43 |
| LC2 | MetaFlow | 15 | 0 | 0 | 1.000 | 1.000 | **1.000** | **0.66** |
|  | MetaPhlAn | 15 | 2 | 0 | 1.000 | 0.882 | 0.938 | 13.74 |
|  | BLAST | 15 | 5 | 0 | 1.000 | 0.750 | 0.857 | 8.33 |
|  | GSMer | 8 | 36 | 7 | 0.533 | 0.182 | 0.271 | N/A |
|  | mOTU | 14 | 3 | 1 | 0.933 | 0.824 | 0.875 | 54.06 |
| LC3 | MetaFlow | 14 | 1 | 1 | 0.933 | 0.933 | 0.933 | 2.12 |
|  | MetaPhlAn | 15 | 1 | 0 | 1.000 | 0.938 | **0.968** | 37.2 |
|  | BLAST | 15 | 2 | 0 | 1.000 | 0.882 | 0.938 | **1.98** |
|  | GSMer | 9 | 45 | 6 | 0.600 | 0.167 | 0.261 | N/A |
|  | mOTU | 14 | 1 | 1 | 0.933 | 0.933 | 0.933 | 5.61 |
| LC4 | MetaFlow | 15 | 0 | 0 | 1.000 | 1.000 | **1.000** | **1.71** |
|  | MetaPhlAn | 15 | 1 | 0 | 1.000 | 0.938 | 0.968 | 11.54 |
|  | BLAST | 15 | 3 | 0 | 1.000 | 0.833 | 0.909 | 3.99 |
|  | GSMer | 8 | 58 | 7 | 0.533 | 0.121 | 0.198 | N/A |
|  | mOTU | 15 | 2 | 0 | 1.000 | 0.882 | 0.938 | 20.29 |
| LC5 | MetaFlow | 15 | 0 | 0 | 1.000 | 1.000 | **1.000** | **1.19** |
|  | MetaPhlAn | 15 | 1 | 0 | 1.000 | 0.938 | 0.968 | 23.04 |
|  | BLAST | 15 | 4 | 0 | 1.000 | 0.789 | 0.882 | 12.2 |
|  | GSMer | 9 | 5 | 6 | 0.600 | 0.643 | 0.621 | N/A |
|  | mOTU | 15 | 1 | 0 | 1.000 | 0.938 | 0.968 | 35.22 |
| LC6 | MetaFlow | 15 | 0 | 0 | 1.000 | 1.000 | **1.000** | 0.48 |
|  | MetaPhlAn | 15 | 0 | 0 | 1.000 | 1.000 | **1.000** | 23.63 |
|  | BLAST | 15 | 0 | 0 | 1.000 | 1.000 | **1.000** | **0.13** |
|  | GSMer | 8 | 13 | 7 | 0.533 | 0.381 | 0.444 | N/A |
|  | mOTU | 15 | 0 | 0 | 1.000 | 1.000 | 1.000 | 2.13 |
| LC7 | MetaFlow | 15 | 0 | 0 | 1.000 | 1.000 | **1.000** | **0.49** |
|  | MetaPhlAn | 15 | 2 | 0 | 1.000 | 0.882 | 0.938 | 18.31 |
|  | BLAST | 15 | 1 | 0 | 1.000 | 0.938 | 0.968 | 0.54 |
|  | GSMer | 8 | 52 | 7 | 0.533 | 0.133 | 0.213 | N/A |
|  | mOTU | 14 | 0 | 1 | 0.933 | 1.000 | 0.966 | 11.84 |
| LC8 | MetaFlow | 15 | 0 | 0 | 1.000 | 1.000 | **1.000** | 1.09 |
|  | MetaPhlAn | 15 | 0 | 0 | 1.000 | 1.000 | **1.000** | 2.76 |
|  | BLAST | 15 | 0 | 0 | 1.000 | 1.000 | **1.000** | **0.7** |
|  | GSMer | 6 | 103 | 9 | 0.400 | 0.055 | 0.097 | N/A |
|  | mOTU | 15 | 0 | 0 | 1.000 | 1.000 | 1.000 | 2.29 |
| LC9 | MetaFlow | 15 | 0 | 0 | 1.000 | 1.000 | **1.000** | 0.49 |
|  | MetaPhlAn | 15 | 1 | 0 | 1.000 | 0.938 | 0.968 | 3.2 |
|  | BLAST | 15 | 0 | 0 | 1.000 | 1.000 | **1.000** | **0.15** |
|  | GSMer | 13 | 20 | 2 | 0.867 | 0.394 | 0.542 | N/A |
|  | mOTU | 14 | 1 | 1 | 0.933 | 0.933 | 0.933 | 27.46 |
| LC10 | MetaFlow | 15 | 0 | 0 | 1.000 | 1.000 | **1.000** | **0.88** |
|  | MetaPhlAn | 15 | 0 | 0 | 1.000 | 1.000 | **1.000** | 16.74 |
|  | BLAST | 15 | 3 | 0 | 1.000 | 0.833 | 0.909 | 1.88 |
|  | GSMer | 11 | 91 | 4 | 0.733 | 0.108 | 0.188 | N/A |
|  | mOTU | 15 | 1 | 0 | 1.000 | 0.938 | 0.968 | 17.45 |
| LC11 | MetaFlow | 15 | 1 | 0 | 1.000 | 0.938 | **0.968** | **9.74** |
|  | MetaPhlAn | 15 | 1 | 0 | 1.000 | 0.938 | **0.968** | 56.09 |
|  | BLAST | 15 | 7 | 0 | 1.000 | 0.682 | 0.811 | 17.48 |
|  | GSMer | 11 | 70 | 4 | 0.733 | 0.136 | 0.229 | N/A |
|  | mOTU | 13 | 2 | 2 | 0.867 | 0.867 | 0.867 | 56.61 |
| LC12 | MetaFlow | 15 | 2 | 0 | 1.000 | 0.882 | **0.938** | **13.9** |
|  | MetaPhlAn | 14 | 4 | 1 | 0.933 | 0.778 | 0.848 | 26.77 |
|  | BLAST | 15 | 5 | 0 | 1.000 | 0.750 | 0.857 | 20.78 |
|  | GSMer | 9 | 81 | 6 | 0.600 | 0.100 | 0.171 | N/A |
|  | mOTU | 12 | 1 | 3 | 0.800 | 0.923 | 0.857 | 98.17 |
| LC13 | MetaFlow | 15 | 3 | 0 | 1.000 | 0.833 | 0.909 | **40.81** |
|  | MetaPhlAn | 14 | 2 | 1 | 0.933 | 0.875 | 0.903 | 56.72 |
|  | BLAST | 15 | 7 | 0 | 1.000 | 0.682 | 0.811 | 41.04 |
|  | GSMer | 8 | 62 | 7 | 0.533 | 0.114 | 0.188 | N/A |
|  | mOTU | 14 | 1 | 1 | 0.933 | 0.933 | **0.933** | 54.81 |
| LC14 | MetaFlow | 15 | 3 | 0 | 1.000 | 0.833 | **0.909** | 60.74 |
|  | MetaPhlAn | 13 | 3 | 2 | 0.867 | 0.813 | 0.839 | 64.94 |
|  | BLAST | 15 | 5 | 0 | 1.000 | 0.750 | 0.857 | **60.4** |
|  | GSMer | 7 | 67 | 8 | 0.467 | 0.095 | 0.157 | N/A |
|  | mOTU | 12 | 1 | 3 | 0.800 | 0.923 | 0.857 | 104.94 |
| LC15 | MetaFlow | 15 | 2 | 0 | 1.000 | 0.882 | **0.938** | 21.19 |
|  | MetaPhlAn | 15 | 2 | 0 | 1.000 | 0.882 | **0.938** | 36.74 |
|  | BLAST | 15 | 3 | 0 | 1.000 | 0.833 | 0.909 | **16.99** |
|  | GSMer | 8 | 102 | 7 | 0.533 | 0.073 | 0.128 | N/A |
|  | mOTU | 10 | 1 | 5 | 0.667 | 0.909 | 0.769 | 46.43 |

**Fig. 3.**
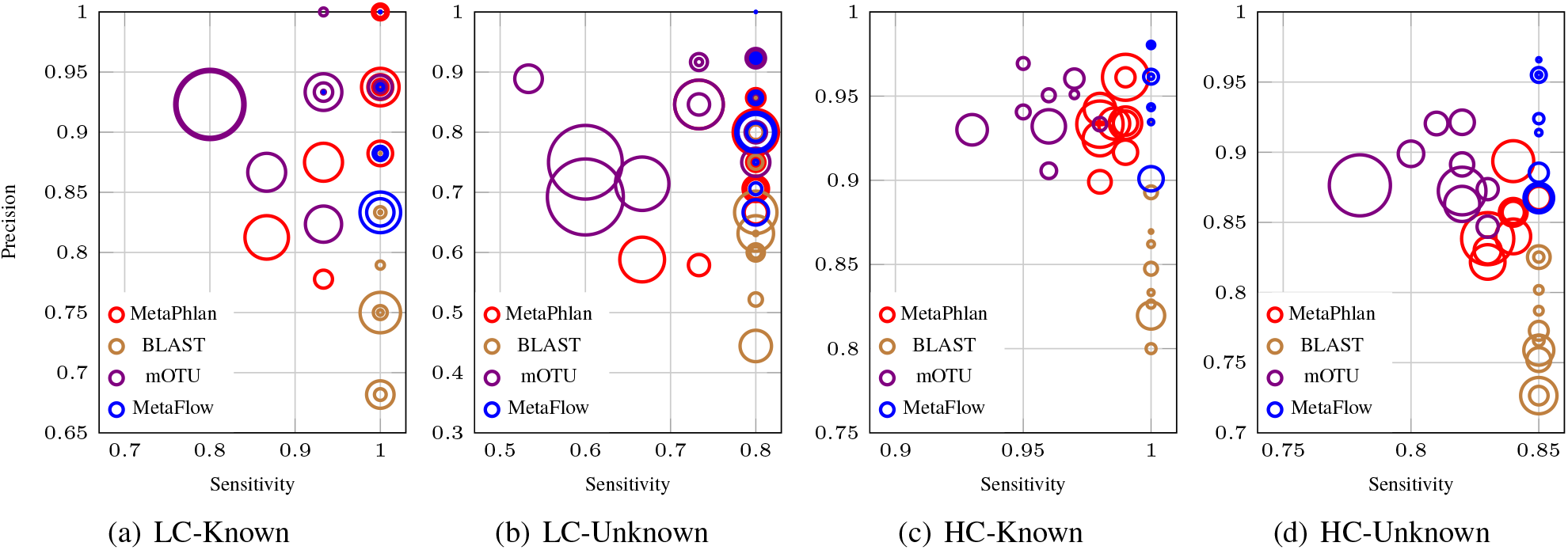
Results of the tools on all simulated data sets. The *x*-axis is the sensitivity and the *y*-axis is the precision; each circle is one experiment; inside each plot, the size of the circles is proportional ‘with the *ℓ*_1_-norm (smaller is better).In LC-Known data set: 15 data sets, each with 4M reads from 1.5 species, all known. In LC-Unknown; 15 data sets, each with 4M reads from 15 is peciea, out ot which 3 are unknown. In HC-Known: 9 data sets, each with 40M reads from 1a0 species, all known. In HC-Unknown: 9 data sets, each with 40M reads from 100 species, out of which 15 are unknown. The unknown spiteies are among 31 bacterial species from the NCBI micsobial genome database, published after 2014. GSMer’s results are not included in the figures, since its precision and sensitivity were always much lower than the other methods (see Table 1).

These results also confirm our hypothesis that taking into account the coverage across the entire genome improves the abundance estimation. For example, even though BLAST has a lower F-measure, it has better *ℓ*_1_-norm than marker-based methods. This is due to the fact that marker regions are much shorter compared to the genome length, and thus slight variations in coverage in these regions can easily skew the abundance estimation. MetaFlow achieves the maximum sensitivity of BLAST, but it manages to highly improve the precision, and it also obtains the best F-measure among all the tested methods. Comparing the LC and HC scenarios, we can also observe that MetaFlow’s richness estimations are robust with the increase in sample size. Moreover, since MetaFlow is also based on whole-genome read alignments, it gives a much better abundance estimation than marker-based methods, with an improvement of 2-4 × in average *ℓ*_1_-norm. Our problem formulation also filters out outlier species and false alignments, thus improving the abundance estimations over BLAST.

Finally, the results are always better on the HC data sets than on the LC data sets for all tools, because a small variation is more severe in a sample with 15 species than in a sample with 100 species. Recall also that the LC data sets were constructed to have similar species, whereas in the HC data sets the species were randomly chosen. The data sets with high genotypic homogeneity (LC13-LC15) show that such scenario remains a difficult one: even though MetaFlow improves both the richness and abundance estimation of the competing methods, its precision drops to an average of 0.85 and its *ℓ*_1_-norm increases to an average of 40.

### 4.2 Real data set

On the fecal metagenomic sample (see Fig. 5 and Table 7 in the Supplementary Material), the most abundant species reported by MetaFlow is *B.uniformis* (23.6% relative abundance), which was also reported as the most abundant species in human feces [16]. This high abundance is also supported by the fact that 15,418,699 reads are mapped by BLAST only to *B.uniformis*. Out of these, 10,721,492 are finally assigned by MetaFlow to *B.uniformis*, because of the uneven read coverage. This corresponds to an average read coverage of 220. To put this number into perspective, the 10th most abundant species according to MetaFlow, *A.shahii*, has relative abundance 2.3% and average read coverage 21. MetaPhlAn and mOTU assign to *B.uniformis* abundances 1.7% and 6.4%, respectively.

**Table 6.**
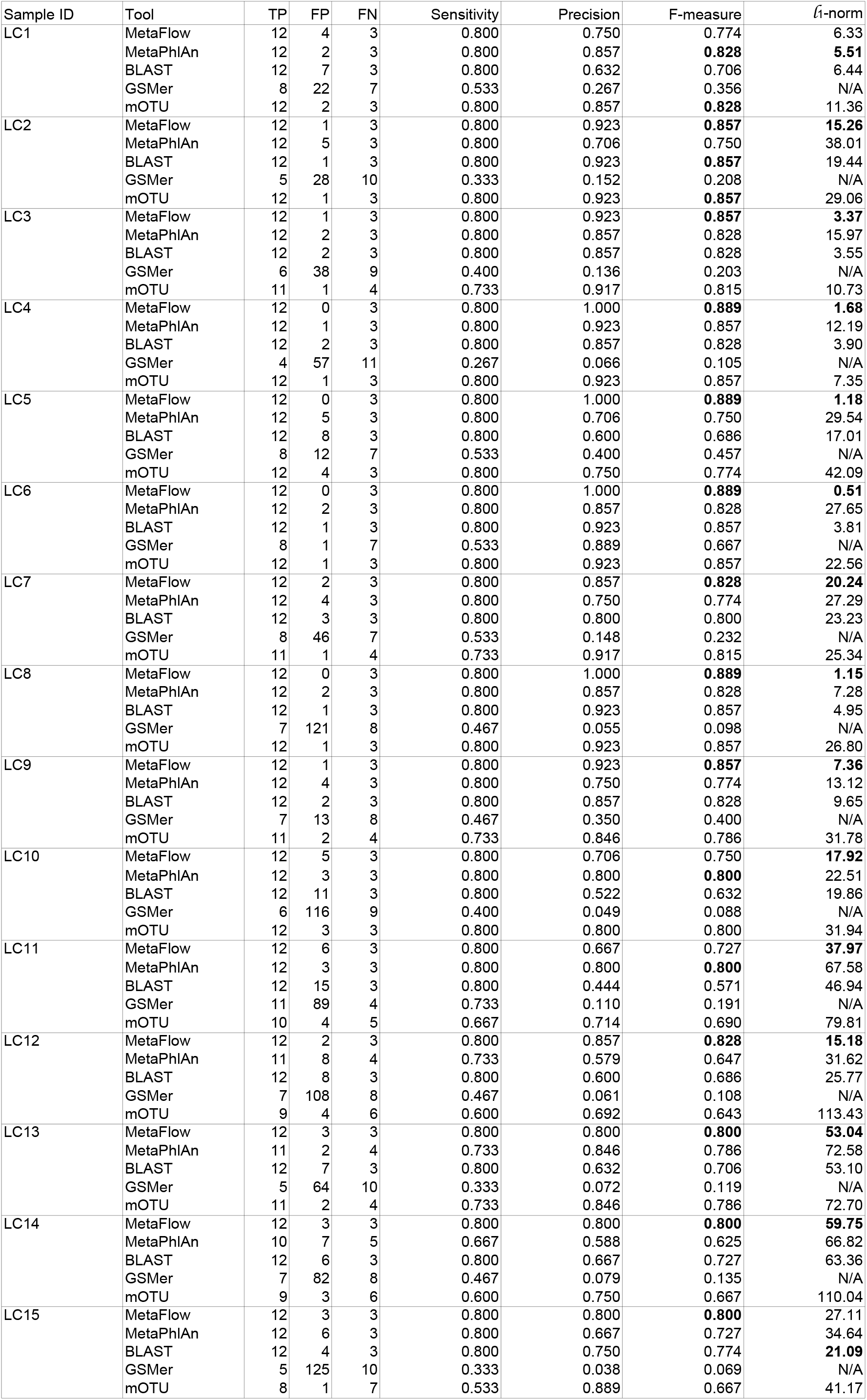
Detailed results on the LC-Unknown data sets. Sample descriptions are in Table 4. Best results are shown in bold. TP = true positives, FP = false positives, FN = false negatives.

The second most abundant species reported by MetaFlow is *B.vulgatus*, another common species in human feces [16]. MetaFlow’s predicted abundance is 22.3% (average read coverage 208), which is in line with MetaPhlAn’s prediction of 17.7% and, to an extent, mOTU’s prediction of 11.9%. In Table 7 of the Supplementary Material we give the list of the top 10 prediction of MetaFlow, and their abundances reported by MetaPhlAn and mOTU. Four species out of the top six species have also been reported by [16] as predominant in human feces, and they constitute 59% of the sample according to MetaFlow (relative to the species known to MetaFlow in this experiment).

**Table 7.**
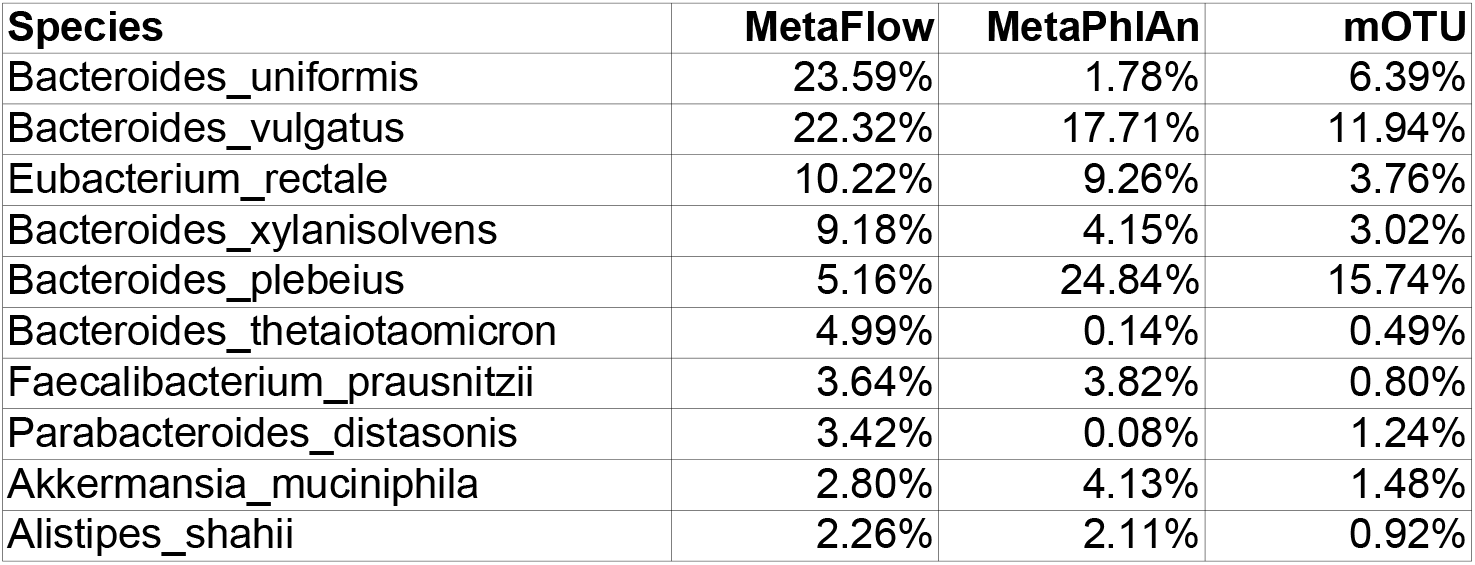
The top 10 species reported by MetaFlow on the real metagenomic sample. *B.uniformis* has been found as the most abundant species in human feces [16], with *B.vulgatus, B.xylanisolvens, F.prausnitzii* also being commonly reported [16].

The top abundant species in MetaPhlAn’s and mOTU’s predictions is *B.plebeius*, with 25% and 16% relative abundance, respectively. MetaFlow’s reported abundance is 5.2%. Note also that 62 species reported by MetaPhlAn are not available in the database of reference genomes given to MetaFlow (NCBI’s database plus *B.uniformis* and *B.plebeius*). Since our predicted abundances are relative to the known genomes only (average read coverages are also outputted by MetaFlow), the abundance of *B.uniformis* relative to all species in the sample may be lower than 23.6%, but it cannot be significantly lower than the one of *B.vulgatus*, as MetaPhlAn and mOTU predict.

## 5 Conclusion

Analysis of high-throughput sequencing samples of metagenomes is crucial for understanding diseases and the environment. This involves discovering what bacterial species are present in a sample, and what are their relative frequencies. State-of-the-art methods are based only on marker regions of known genomes, which can heavily skew the abundances, especially at the species level.

MetaFlow is the first method based on coverage analysis across entire known genomes which also scales to HTS samples. MetaFlow shows consistent results across a wide range of synthetic data sets, with abundance estimations two to four times better, in terms of *ℓ*_1_-norm, than popular tools. On a real human stool data set MetaFlow also proves robust, whereas marker-based methods report highly skewed results. Finally, our whole-genome coverage method makes obsolete the need to curate and continuously update a database of genomic markers.

## Acknowledgement

We thank Romeo Rizzi for discussions about the computational complexity of our problem formulation. This work was partially supported by the Academy of Finland under grants 284598 (CoECGR) to A.S. and V.M. and 274977 to A.T.

## Footnotes

A shorter version of this manuscript will appear in the Proceedings of RECOMB 2016, 20th Annual International Conference on Research in Computational Molecular Biology.

## Supplementary Material

### A Methods

#### A.1 Definitions of flows

We now recall some standard definitions about flows, following [12].

**Definition 1**. *A* flow network *is a tuple N* = (*G, ℓ, u, c, q*), *where*

– *G is a directed graph (with unique source s and unique sink t);*
– *ℓ and u are functions assigning a non-negative integer* demand *and* capacity, *respectively, to every arc;*
– *c is a function assigning a* cost per unit of flow *to every arc;*
– *q is the required* value *of the flow*.

A flow over a flow network is a function satisfying the flow conservation property, the demand and capacity constraints. The *value* of a flow is the sum of the flow exiting the source of the flow network. More formally:

**Definition 2**. *Given a flow network N* = (*G, ℓ, u, c, q*), *a function f: E(G)* → ℕ *is a* flow *over the network N if the following conditions hold:*

– **Flow conservation**: *for every vertex x ∈ V(G) \ {s,t}, it holds that*

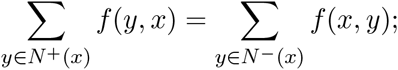
– **Demand and capacity constraints**: *for every arc (x,y) ∈ E(G), it holds that*

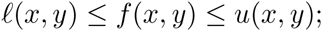
– **Required flow value** *q*: 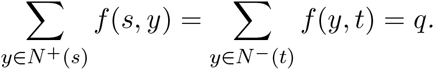

**Definition 3**. *A flow f over a flow network N* = (*G,ℓ,u,c,q*) *is called a* min-cost flow *if, among all flows over N, it minimizes*

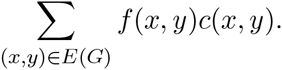

See e.g., [12] for more details on min-cost flows. We used the LEMON library [3], which is an open source C++ library, which provides implementations of different data structures and algorithms. It provides an implementation of the network simplex algorithm [8] which we used for solving our min-cost flow problem.

### B Further implementation details

#### B.1 Input

MetaFlow was written in C++, and it runs from the command line. There are two input files for the tool:

**A mapping file:** This is the aligner’s output file converted into another format which is easier to process by MetaFlow. We provide a Python script for converting BLAST output into our mapping file. If the user wishes to use another aligner, she should convert the aligner’s output into our mapping file format.
**A species database file:** A file containing the names and lengths of the genomes in the reference database. We provide a default file created from NCBI bacterial database. If another reference database will be used, a species database file should be created from the new database.

MetaFlow has an online tutorial that explains the format of these files and how to create them.

#### B.2 Output

MetaFlow generates different output files. All the files are in CSV format to make any further analysis easy. Table 3 lists the output files and their description.

#### B.3 metaflow.config configuration file

We provide a set of configuration parameters which can be used to tune the behavior of our model for finding the approximated relative abundance, and to control the running time of MetaFlow. The metaflow.config is a configuration file that contains these parameters. The parameters are new line-separated, whereas the parameters and their values are tab-separated. Here, we list these parameters, their description and default values.

**CHUNK_SIZE:** Each genome is split into chunks of equal sizes, to which the reads are mapped based on their mapping position given by the aligner. If the chunk size is too large, e.g. 50,000, then the uniformity assumption in read coverage is not fully exploited. On the other hand, if it is too short, e.g. 500, then the running time becomes unfeasible. The chunk size should be decided based on the average read length. For example, it can be selected to be ten times the average read length (default=2000).
**ALPHA** (*α*): The weight of the uniform distribution constraint in the optimization problem explained in the main paper. If *α* is too small, the aligner mapping score will dominate, and the uniform distribution constraint will be almost neglected. If *α* is too large, uniform distribution constraint will dominate, and the mapping score will be almost neglected (*α* ∈ [0,1], default=0.9).
**TRIMMING_PERCENTAGE** (*TR*): As explained in the main paper, we iterate the min-cost flow algorithm, at each step updating the solution vector *R^j^* to *R*^*j*+1^, using the *TR*-trimmed mean read coverage of the chunks of each genome *G^i^*, obtained from *M^j^. TR* is thus the trimmed mean value (0 ≤ *TR* ≤ 0.5, default=0.2). A trimmed mean with value *TR* = 0.2 is calculated by trimming 40% of the chunks (the lowest 20% and highest 20% based on their coverage), and calculating the mean for the rest 60%. If *TR* = 0, then the required abundance per chunk in *G^i^* will be the mean coverage for all chunks. If *TR* = 0.5, the required abundance per chunk will be the median coverage of the chunks.

In practice, it is not true that the read coverage of each base (in our case, of each chunk) is the same across each reference genome, but it follows a distribution. The are various proposals for this distribution, for example the negative binomial [14] or gamma [6], or mixtures of distributions [10]. In order to achieve a better approximation of the true distribution, we introduce two parameters for relaxing the objective function:

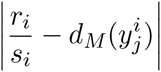

of the coverage-sensitive metagenomic mapping problem and allowing for a more practical formulation. The two are the following:

**SPLIT_ARCS** (*SA*): A boolean value (0 for false and 1 for true) allows for switching between a strict and a relaxed uniform distribution constraint. If *SA* = 1, the uniform distribution constraint is relaxed, allowing chunks to have coverage higher or lower than the required coverage based on the value of the parameter NUM_OF_READS_WITH_LOWER_COST (see below). This is achieved by penalizing these variations less than the default coverage penalty. If *SA* = 0 any variations from the required coverage will be penalized with the default penalty, which makes the uniform distribution constraint strict (default=1).
**NUM_OF_READS_WITH_LOWER_COST** (*N_lc_*): The number of reads per each chunk that will be penalized less in case of coverage lower or higher than the required one (*N_lc_* ∈ *N*^0^, default=5). Fig. 4 gives an example of how these two parameters relax the uniform distribution constraint.

**Fig. 4.**
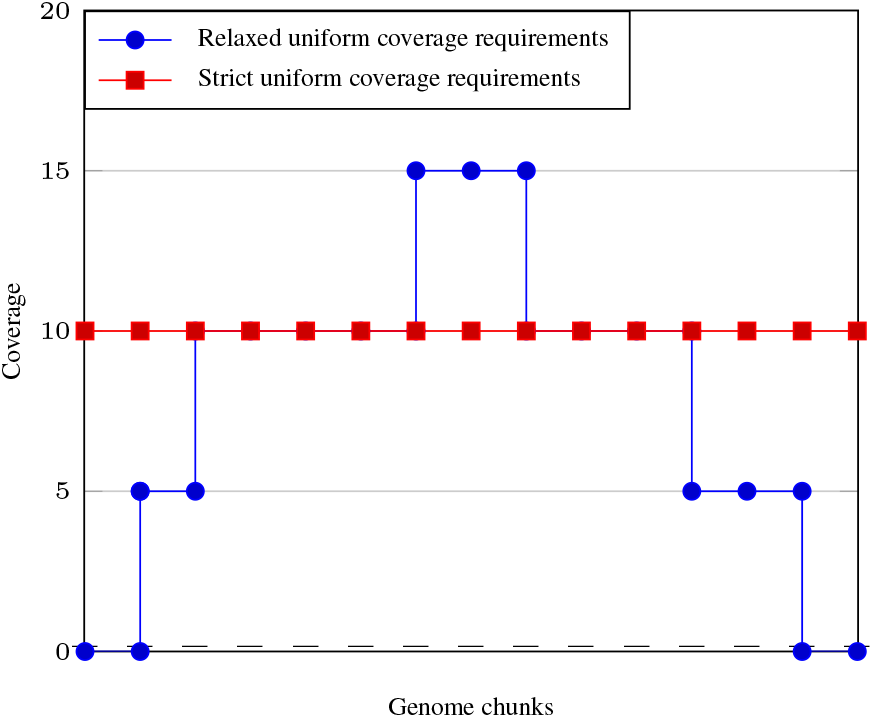
Ctamparison between the strist and relaxed uniform distribution requirments. The strict uniform distribiution requimient (in rem) requires all cliunk;s to be covered with 10 reads. Whe relaxed uniform distri bution requirement (in blue) also requires all chunks to be covered with 10 reads. However, the parameter NUM_OF_READS_WITH_LOWER_COST=5 allows for some chunks to have coverage 5, and some other-chunks for have coverage up to 15. Also, the parameter REQUIRED_MAX_PER_OF_EMPTY_CHUNKS allows for having completely empty chunks. This allows for a better approximation for tge true distribution of the leads.

**Fig. 5.**
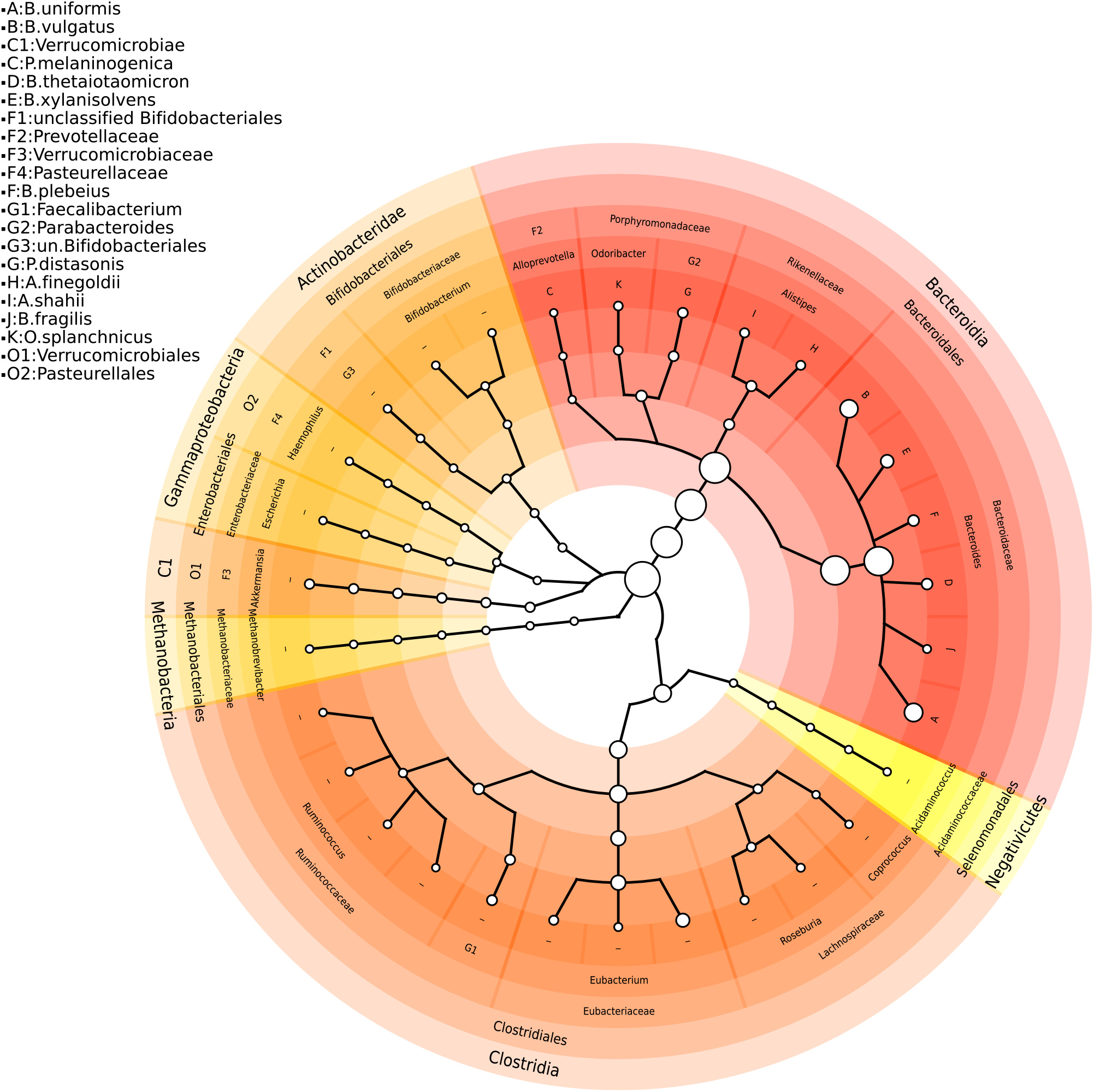
The species found by MetaFlow in the real sample. This sample was obtained by merging 6 G_DNA_Stool samples from the Human Microbiome Project (287M reads out of which 98.2M BLAST mapped to some species whose reference was available to Metaflow). The size of the circles is proportional to the relative abundance at that taxonomic level.

This idea is similar to the one explained in e.g., [12, pp. 51-52], on solving min-cost flows with convex cost functions.

Deciding whether a species is an outlier or not is based on the absolute abundance of the species, and on the coverage of its chunks. For example, if the absolute abundance of a species is 0.1, then with high probability it is an outlier, and should be removed from the sample. We use the following parameters for deciding whether a species is an outlier or not. The user can choose to relax or tighten their values based on her knowledge about the input same.

**REQUIRED_MIN_ABUNDANCE** (*rel_abundance_min_*): For any species, if the absolute abundance is lower than *rel_abundance_min_*, then the species is considered an outlier and will be removed (*rel_abundance_min_* ∈ [0,1], default=0.3).
**REQUIRED_AVERAGE_CHUNKS_COVERAGE** (*Coverage_avg_*): For any species, if the average coverage of chunks (number of reads/number of chunks) is lower than *Coverage_avg_*, then the species is considered an outlier and will be removed (*Coverage_avg_* ∈ ℕ, default=2).
**REQUIRED_MAX_PER_OF_EMPTY_CHUNKS** (*Ckunks_emp_*): For any species, if the ratio between number of chunks not covered by any read and the total number of chunks is more than *Chunks_emp_*, than the species will be considered an outlier and will be removed (*Chunks_emp_* ∈ [0,1], defauft=0.4).

The size of the flow netwonk (numbee of nodes, and number of arcs) affects the runnimg time for the MCF solver. (The following parameters are used for honfoolling the number of arcu in a flow network.

**MAX_SCORE_DIFFERENCE** (*S_diff_*): For each read, if the difference between the score of the best hit and any other hit is larger than *S_diff_*, remove thin hit (*S_diff_* ∈ ℕ, default=0).
**MAX_NUMBER_OF_ARCS**: If the number of arcs in a flow network is larger than this value, randomly remove arcs with higher mapping cost until the number of arcs is less than this (default=5,000,000).

As explained before, finding an approximation for the optimum abundances is achieved by running the min-cost flow algorithm iteratively. Tire following parameters are used for controlling the running time.

**MAX_COST_DIFFERENCE** (*C_diff_*): For any two consecutive iterations *itr_k_, itr*_*k*+1_ with solution costs *C_k_,C*_*k*+1_, respectively if |*C_k_* – *C*_*k*+1_| ≤ *C_diff_*, then termmate the loop cnd return the solution of *itr*_*k*+1_ approximated aleundances (*C_diff_* ∈ ℕ, default=1).
**MAX_READS_DIFFERENCE** (*R_diff_*): For any two consecutive if tire difference between the total number of reads remaining after each deration is larger than *R_diff_*, terminate the loop and neturn the current solutfon (*R_diff_* ∈ ℕ, default=0).
**MAX_NUMBER_OF_LOOPS** (*N_loops_*): The total number of iterationf for finding the approximated abundance. If the number of iterations exceeded this value, terminate the loop and return the current solution as the approximated abundance (*N_loops_* ∈ ℕ^+^, default=10).
**MAX_RUNNING_TIME**: If the running time (in minutes) for finding the approximated abundance exceeded this value, terminate the loop and return the current solution as the approximated abundance (default=300).

### C Further experimental details

The complete results of the simulated low complexity samples are in Tables 5, 6. The real samples used were downloaded from NCBI SRA database with accession numbers (SRX877403, SRX877395, SRX877412, SRX024197, SRX025210, SRX877173). The reads from the pyrosequencing sample were split into reads of length 100bp.

